# Haplo-insufficiency of Profilin1 in vascular endothelial cells is beneficial but not sufficient to confer protection against experimentally induced atherosclerosis

**DOI:** 10.1101/2023.12.06.570450

**Authors:** Abigail Allen-Gondringer, David Gau, Partha Dutta, Partha Roy

## Abstract

Actin cytoskeleton plays an important role in various aspects of atherosclerosis, a key driver of ischemic heart disease. Actin-binding protein Profilin1 (Pfn1) is overexpressed in atherosclerotic plaques in human disease, and Pfn1, when partially depleted globally in all cell types, confers atheroprotection *in vivo*. This study investigates the impact of endothelial cell (EC)-specific partial loss of Pfn1 expression in atherosclerosis development. We utilized mice engineered for conditional heterozygous knockout of the Pfn1 gene in ECs, with atherosclerosis induced by depletion of hepatic LDL receptor by gene delivery of PCSK9 combined with high-cholesterol diet. Our studies show that partial depletion of EC Pfn1 has certain beneficial effects marked by dampening of select pro-atherogenic cytokines (CXCL10 and IL7) with concomitant reduction in cytotoxic T cell abundance but is not sufficient to reduce hyperlipidemia and confer atheroprotection *in vivo*. In light of these findings, we conclude that atheroprotective phenotype conferred by global Pfn1 haplo-insufficiency requires contributions of additional cell types that are relevant for atherosclerosis progression.

## Introduction

Cardiovascular diseases are a leading cause of death world-wide, contributing to approximately 17.9 million deaths a year (1). The most common form, ischemic heart disease, remains the leading cause of cardiovascular-related death, and develops due to formation of atherosclerotic plaques in the blood vessels (2). Atherosclerosis begins with fatty streak deposition, appearing in clinically relevant locations including aortic, coronary, and cerebellar arteries as early as age 30-40 years in humans. Over time, fatty streaks develop into fibrotic plaques composed of extracellular matrix and necrotic cellular deposits, smooth muscle cells (SMCs) and immune cells. Immune cell sub-populations are characterized by infiltration of lipid-rich macrophagic foam cells, cytotoxic T lymphocytes, granulocytes, and other antigen presenting cells (APCs) (3,4). An early contributor to fatty streaks and plaque development is EC dysfunction. In the healthy endothelium, atheroprotective nitric oxide (NO) produced by the action of endothelial nitric oxide synthase (eNOS) regulates the expressions of various endothelial cell adhesion molecules (CAMs, i.e. V(vascular)CAM-1, I(intracellular)CAM-1) (5). Endothelial NOS dysfunction in the vasculature (signified by a decrease in bioavailable nitric oxide (NO)) enhances the levels of VCAM-1 and ICAM-1, inciting local immune invasion and inflammation(6) (7).

Actin cytoskeleton plays an important role in various aspects of atherosclerosis. For example, remodeling of actin cytoskeleton is largely responsible for pro-atherogenic morphological changes and dysfunction (characterized by impaired barrier function) of ECs in response to turbulent flow (8–10). Actin regulatory proteins control EC stiffness, in turn impacting clustering of cell surface adhesion molecules and transendothelial migration of immune cells (11). Other aspects of atherosclerosis, such as polarization and lipid processing by macrophages, are also sensitive to the state of actin cytoskeleton in immune cells (12,13). In human atherosclerosis, actin-binding protein Profilin1 (Pfn1 - an important promoter of actin polymerization in cells) is overexpressed in atherosclerotic plaques and present at a higher circulating level in the serum (14). High-fat diet-induced atherosclerosis in low-density lipoprotein receptor (LDLR) knockout (LDLR^−/−^) mice also leads to elevated Pfn1 expression increase in atherosclerotic blood vessels mimicking the human disease (15). Furthermore, when Pfn1 expression is partially depleted globally in all cell types by heterozygous gene deletion in LDLR^−/−^ mice, it results in reduced severity of atherosclerosis marked by smaller plaque size, lower VCAM-1 expression, and diminished vascular infiltration by macrophages (16). However, given that atherosclerosis initiation and progression involves interplay between a number of different cell types including vascular EC, immune cells (monocytes/macrophages, neutrophils and various subpopulation of T cells) and vascular smooth muscle cells, Pfn1 downregulation in which specific cell type is primarily responsible for atheroprotective effect in Pfn1^+/−^ mice remained unclear. In the present study, we sought to investigate whether haplo-insufficiency of Pfn1 in vascular ECs alone has any beneficial effect in experimentally induced atherosclerosis.

## Material and Methods

### *In vivo* experiments

Pfn1^flox/flox^:CDH5-Cre-ERT2 mice, as previously described (17,18) were backcrossed into C57Bl/6 background for 5-6 generations to generate Pfn1^flox/WT^:CDH5-Cre-ERT2 mice (WT – wild-type allele). These mice (both male and female) when 4-6 weeks old, were subjected to intraperitoneal injection of 100μl tamoxifen (dissolved in peanut oil at a concentration of 10 mg/ml) daily over a course of five days to achieve Cre-mediated deletion of the floxed Pfn1 allele (denoted as ^Pfn1+/−(EC)^ hereon), as we had done before(17). Control animals (any Pfn1 genotype without Cre expression - denoted as Pfn1^+/+^ hereon) were also subjected to tamoxifen administration in a similar manner (17). Details of genotyping primers to confirm floxed alleles of Pfn1, Cre, and Cre-mediated excision of Pfn1 are described in (19). To confirm Cre-mediated downregulation of Pfn1 expression at the protein level, CD31+ ECs were isolated from the kidneys (a vascular-rich organ) of select Pfn1^+/+^ and Pfn1^+/−(EC)^ mice, and cultured for 7 days before performing immunoblot analyses of total cell extracts as per our previously described protocol (17). One month after gene excision, mice were subjected to intraperitoneal injection of 2×10^11^ PFU AAV8 encoding constitutively active mutant (D377Y) form of mouse PCSK9 (Proprotein convertase subtilisin/kexin type-9; referred to as AAV8-mPCSK9 from hereon; Vector Biolabs, Malvern PA) and fed a high-cholesterol diet (HCD - D12336i, Research Diets, New Brunswick NJ) for 3 months to develop atherosclerosis (20,21). For confirmation of LDLR knockout and serum cholesterol elevation, animals were similarly injected with GFP-encoding AAV8 as control. Animals were euthanized for whole blood isolation by cardiac puncture and harvesting of various organs of interest (aorta, spleen, and long bone). All animal experiments were performed in compliance with an approved IACUC protocol, according to the University of Pittsburgh Division of Laboratory Animal Resources guidelines.

### Lipid and Cytokine/chemokine Analyses

To probe for cytokine/chemokine expression, mouse serum (prepared from whole blood and further diluted with PBS at a 1:1 ratio) were analyzed by Luminex service provided by Eve Technologies (Calgary, AB Canada) using MD32, a 32-analyte murine discovery panel. Analyte values were normalized to the average of wild-type animals across litters. Lipid panel analyses of mouse serum were performed for measurements of low-density lipoprotein (LDL), high-density lipoprotein (HDL), total cholesterol, and triglycerides utilizing services provided by the University of Pittsburgh Division of Laboratory Animal Resources (Idexx #6290, Westbrook ME) to confirm lipid elevation in atherosclerotic mice.

### Histology and quantification of atherosclerosis parameters

Briefly, cardiac tissue was fixed overnight, maintained in 30% sucrose, and excess cardiac tissue was cut up to the aortic root and cleaned of excess tissue before embedding in OCT for performing 5-10 μm serial sections until the aortic root could be appropriately visualized. These sections were stained for Oil Red-O to visualize lipid accumulation, and Masson’s trichrome to visualize the fibrous cap and collagen deposition. These services were performed by the University of Pittsburgh histology core. Total plaque was identified by dark-red area present in Oil Red O staining. Fibrous cap was identified as darkly stained layer present around necrotic core (situated between vascular wall and fibrous cap). Using ImageJ, histology images were converted to 8-bit grayscale images before selecting the region of interest (ROI) using the polygon tool and further thresholding to compute the areas of the ROIs.

### Immunological profiling

Whole blood or various tissues (spleen, thoracic aorta, bone marrow) were crushed and passed through a 70μm cell strainer, treated with 1X red blood cell (RBC) lysis buffer to reduce RBC contamination, and incubated with Brilliant Stain Buffer (BD Biosciences, San Jose, CA) to prevent signal overlap between the blue and the ultraviolet markers. Samples were stained with either a full panel of immune-cell specific antibodies or buffer only for 30 minutes at 4°C, spun, and fixed in fixation/permeabilization buffer overnight before staining for various immunological markers for 30 minutes at room temperature (see (17) for details of immune-panel antibodies and respective dilution for staining). Samples were run on a Fortessa FACS Aria II and the data were analyzed by FlowJo (FlowJo Inc., Ashland, OR) software.

### Immunoblot

Total cell lysate (TCL) was collected using a modified RIPA buffer (25 mMTris-HCl, pH 7.5, 150 mM NaCl, 1% (v/v) Nonidet P-40, 5% (v/v) glycerol), 1 mM EDTA, 50 mM NaF, 1 mM sodium pervanadate, along with 6x sample buffer with SDS diluted to 2% in final buffer. For liver extract preparation, mouse liver was cut, digested, and sonicated in lysis buffer for 1 min and spun for 30 min at >8000g. For immunoblotting, antibodies specific for LDLR (BioLegend, ab124904; 1:200), GFP (Living Colors, Takara Biotech, San Jose CA, 632375; 1:2000), Pfn1 (Abcam, ab124904; 1:3000)), and GAPDH (Sigma-Aldrich, G9545; 1:2000) were used.

### F-actin/microtubule staining

Mouse kidney ECs isolated from Pfn1^+/+^ and Pfn1^+/−(EC)^ mice as described earlier were cultured on glass coverslips coated with 10 µg/ml collagen-1. Cells were fixed for 15 min by directly adding 7.4% paraformaldehyde (PFA) in DPBS to the cell culture media to a final concentration of 3.7% PFA. Following permeabilization with 1% triton X-100, (5 min) and blocking with 10% goat serum for 30 min, cells were stained for microtubules by 1 hr incubation with a primary antibody against α-tubulin (Sigma-Aldrich, T5168, 1:200). Following three washes, cells were incubated with FITC-conjugated phalloidin (Thermo Fisher Scentific, F432), and a TRITC-conjugated anti-mouse secondary antibody (Jackson ImmunoResearch, 115-025-003, 1:100) for an additional 1 hr. Stained cells were washed 3X and mounted on a slide with DAPI in the mounting media (Thermo Fisher Scientific, P36935) prior to imaging on an Olympus IX83 wide-field microscope using a 60X, 1.4 N.A. objective.

### Statistics

All statistical tests were performed using GraphPad Prism 9 software. For experiments with small sample size and non-normal distribution of data, non-parametric Mann-Whitney u-test was used for statistical comparison of data between the groups. For experiments involving a larger sample size and normal sample distribution, an unpaired t-test was used. Two-way ANOVA was used for multiple group comparison in lipid panel analysis. A p-value less than 0.05 was considered to be statistically significant.

## Results and Discussion

To assess the impact of EC-specific Pfn1 depletion on atherosclerosis, we used Pfn1^+/−(EC)^ C57Bl/6 mice for two reasons. First, we recently showed that complete loss of endothelial Pfn1 leads to mortality within 3 weeks of gene ablation(17). Since it takes several months to develop atherosclerosis in mouse models, Pfn1^−/−(EC)^ mice are not suitable for atherosclerosis development studies. Second, use of Pfn1^+/−(EC)^ animals (which are completely viable) allows for a direct evaluation of the impact of EC-specific partial depletion of Pfn1 against previously reported phenotype of global Pfn1 heterozygous knockout mouse in an atherosclerosis setting (16). As a part of characterization of ECs from each of the two genotype of mice (Pfn1^+/+^ vs Pfn1^+/−(EC)^), we first performed Pfn1 immunoblot analyses of total extracts of cultured CD31+ kidney ECs harvested from these mice. Based on these analyses, ECs harvested from Pfn1^+/−(EC)^ mice exhibited slightly more than 50% downregulation of Pfn1 expression at the protein level relative to those harvested from Pfn1^+/+^ mice (**Figs 1A-B**). From phase-contrast images, these two groups of ECs were found to be morphologically indistinguishable (**Fig 1C**). To further determine whether haplo-insufficiency of Pfn1 has any discernible impact on either actin and/or microtubule cytoskeleton in ECs, we performed phalloidin and α-tubulin staining. We found that partial loss of Pfn1 expression led to minor reduction in the overall microtubule (by ∼10%) and F-actin (by ∼25%) content in EC (**Figs 1D-F**). However, when we quantified the ratio of area-to-perimeter (a morphological parameter) on a cell-by-cell basis, there was no statistically significant difference between the two groups (**Fig 1G**) - this finding was in agreement with our qualitative morphological assessment of cells from phase-contrast images.

**Figure 1.**
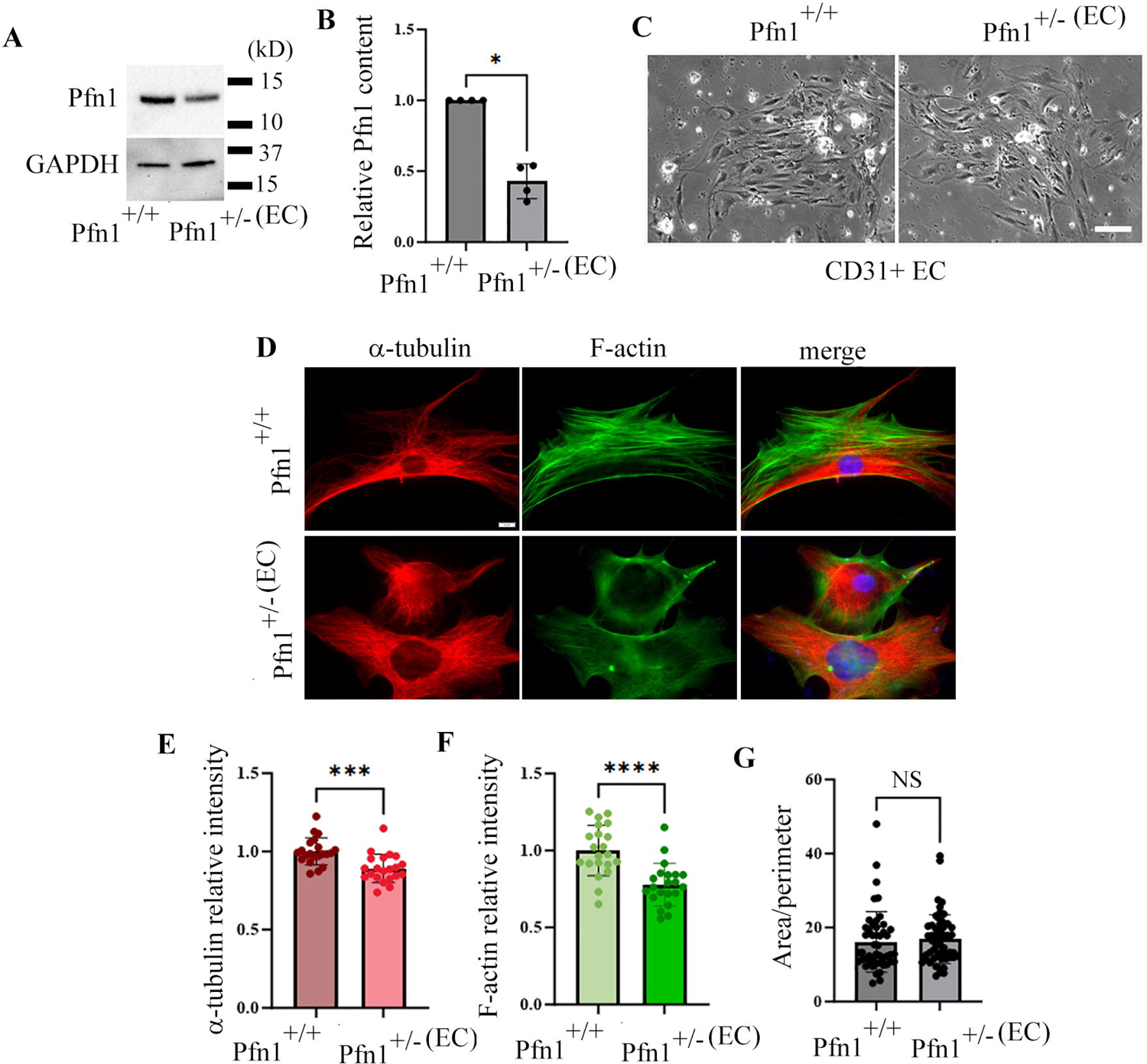
Characterization of vascular ECs with partial loss of Pfn1. **A-B)** Representative Pfn1 immunoblot (*panel A* – GAPDH: loading control) of total cell lysates of CD31+ ECs isolated from Pfn1^+/+^ and Pfn1^+/−(EC)^ mice; panel B summarizes Pfn1 expression in ECs isolated from Pfn1^+/−(EC)^ mice relative to those isolated from Pfn1^+/+^ mice (N = 4 experiments, * p-value < 0.05). **C)** Representative phase-contrast images of CD31+ mouse kidney ECs isolated from Pfn1^+/+^ and Pfn1^+/−(EC)^ mice (scale bar –100 µm). **D-G)** Representative 60x images of CD31+ mouse kidney ECs isolated from Pfn1^+/+^ and Pfn1^+/−(EC)^ mice and stained for microtubules (by α-tubulin antibody), F-actin (by phallodin), and nuceli (by DAPI) (*panel D;* scale bar – 10 μm). Panels E and F summarize the avergae fluorescent intensities of α-tubulin and phalloidin, respectively, in Pfn1^+/−^ relative to Pfn1^+/+^ ECs. Panel G summarzies the average area-to-perimeter ratio of Pfn1^+/−^ ECs relative to Pfn1^+/+^ ECs. These data are based on the analyses of 3 independent experiments (up to 20 cells/group were analyzed per experiment). *** p-value < 0.001, **** p-value < 0.0001, ns – not significant.

Since generating LDLR^−/−^ mice harboring additional cell-type-specific conditional knockout of gene of interest requires extensive breeding and is time consuming, we adopted use of AAV8-mPCSK9 combined with HCD to induce atherosclerosis (20,21). PCSK9 directly binds to hepatic LDLR promoting its degradation in the lysosomes, and consequently, induces hyperlipidemia (22). To initially confirm the efficacy of AAV8-mPCSK9–induced depletion of hepatic LDLR, we probed for expression of LDLR in the liver in both Pfn1^+/+^ and Pfn1^+/−(EC)^ mice 3.5 months after a single intraperitoneal injection of AAV8-mPCSK9 vs AAV8-GFP (as control). Immunoblot analyses of liver extracts showed near complete depletion of LDLR in both Pfn1^+/+^ and Pfn1^+/−(EC)^ mice following administration of AAV8-mPCSK9 (**Fig 2A**; **Fig S1** summarizes the immunoblot data demonstrating an average 84% reduction in hepatic LDLR expression in AAV8-mPCSK9 injected mice relative to those administered with control AAV8-GFP). Lipid panel analyses of serum showed significant elevation of both total cholesterol and LDL (but not triglycerides and HDL) in HCD-fed Pfn1^+/+^ and Pfn1^+/−(EC)^ mice which were administered with AAV8-mPCSK9 relative to those receiving AAV8-GFP (**Fig 2B**). However, we did not find any significant difference in the serum level of either total cholesterol or LDL between Pfn1^+/+^ and Pfn1^+/−(EC)^ animals (**Fig 2B**) suggesting that partial loss of EC Pfn1 does not impact the severity of hyperlipidemia. Oil Red-O staining of aortic roots revealed presence of atherosclerotic plaques in both Pfn1^+/+^ and Pfn1^+/−(EC)^ mice (**Fig 2C**). However, when we compared multiple features of atherosclerosis including the areas of total plaque, fibrous cap and necrotic core measured at the aortic root, we did not find any statistically significant difference in any of these parameters between the two Pfn1 genotypes of mice (**Fig 2D**). These data further suggest that partial depletion of Pfn1 in vascular EC alone is not sufficient to confer overall protection against experimentally induced atherosclerosis *in vivo*.

**Figure 2:**
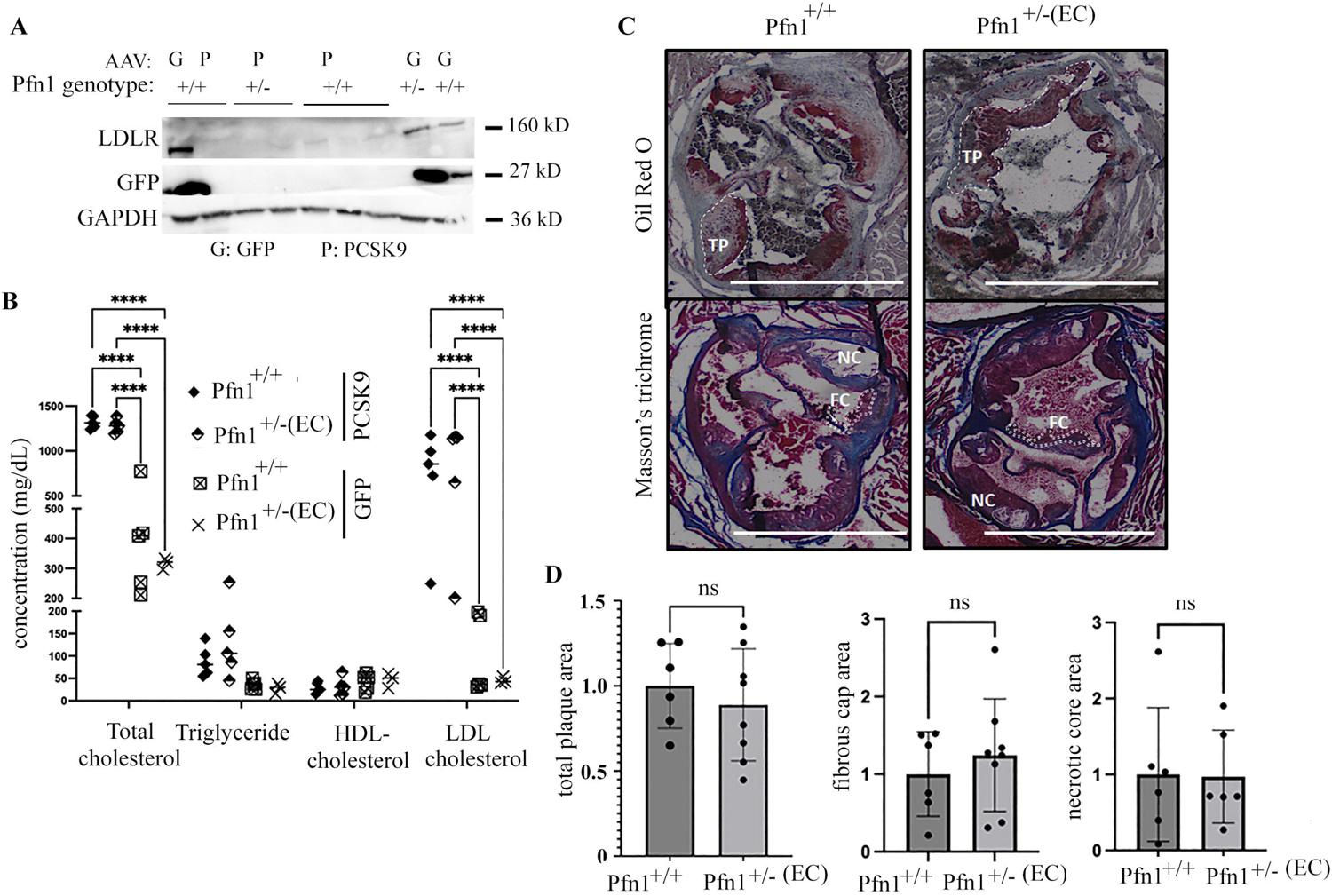
Pfn1 haplo-insufficiency in vascular EC is not sufficient to induce atheroprotection. **A)** Representative immunoblot anlsyses of liver extracts prepared from mice of indicated Pfn1 genotypes showing loss of LDLR expression in reponse to AAV8-mPCSK9 administration (samples from AAV8-GFP injected mice serve as control; GAPDH-loading control). **B)** Relative serum levels of total cholesterol, LDL, HDL and triglycerides in AAV8-mPCSK9 vs AAV8-GFP injected mice of indicated Pfn1 genotypes maintained on a high cholesterol diet. The number of animals for various groups for lipid panel analyses were: Pfn1^+/+^/PCSK9 (n=5); Pfn1^+/−(EC)^/PCSK9 (n=5); Pfn1^+/+^/GFP (n=5); Pfn1^+/−(EC)^/GFP (n=3). **C-D)** Representative images of aortic roots prepared from atherosclerotic Pfn1^+/+^ and Pfn1^+/−(EC)^ mice were stained with Oil Red-O to visualize total plaque (TP) and Masson’s trichrome staining to visualize fibrous cap (FC) and necrotic core (NC) (*panel C*) along with quantifications of various atherosclerosis-related parameters (data normalized to the mean values of Pfn1^+/+^ mice (*panel D*). Data were summarized from 6 Pfn1^+/+^ and 8 Pfn1^+/−(EC)^ atherosclerotic mice pooled from multiple experiments; *p<0.05, **p<0.01, ***p<0.001, ****p<0.0001; ns – not significant.

Vascular infiltration of both innate and adaptive immune cells is a key feature of atherosclerosis. We recently showed that triggering complete loss of vascular EC Pfn1 has a profound immunological consequence providing evidence for Pfn1 as an important mediator of vascular EC-immune cell crosstalk (17). Specifically, mice completely deficient in EC Pfn1 exhibit systemic elevation of pro-inflammatory cytokines and a selective bias toward pro-inflammatory myeloid-derived population of immune cells. Pro-inflammatory M1 macrophages exacerbates the disease progression (23,24). The role of cytotoxic CD8+ T cells in atherosclerosis has been debatable in the literature. CD8+ T cell activity induces growth and promote instability (i.e. rupture-prone) of atherosclerotic lesions through lysis of EC and smooth muscle cells (25). These findings correlate with increased abundance of CD8+ T cells in advanced atherosclerotic lesions (26), and higher circulatory level of CD8+ T cells linked to atherosclerosis development in humans (27). These results support a pro-atherosclerotic role of CD8+ T cells. However, an inverse correlation between CD8+ T cell and macrophage content in human atherosclerotic aortas, taken together with preclinical evidence for antibody-mediated depletion of CD8+ T cells resulting in reduced plaque stability and increased necrosis and macrophage content in advanced-stage LDLR^−/−^ atherosclerotic animals support a protective role of CD8+ T cells in late-stage atherosclerosis (28). In general, immunosuppressive action of regulatory T cells (Tregs) inhibits atherosclerosis (29). Furthermore, B cell-driven antibody response to oxidized LDL (driver of chronic inflammation), and cooperation between B and T cells have been demonstrated to contribute to athero-protective immunity in ApoE^−/−^ mouse model of atherosclerosis (30). Although our studies showed that partial depletion of endothelial Pfn1 alone does not ameliorate either hyperlipidemia or the severity of atherosclerosis, we asked whether haplo-insufficiency of vascular EC Pfn1 has any favorable immunological consequence in the setting of atherosclerosis or otherwise. Therefore, we performed flow cytometry analysis to obtain immunological profile of spleen, bone marrow, thoracic aorta and whole blood isolated from atherosclerotic Pfn1^+/+^ and Pfn1^+/−(EC)^ mice, and compared the findings from these studies to those obtained in a similar manner for non-atherosclerotic mice (these animals did not receive AAV8-mPCSK9 and were maintained in a normal diet). As summarized in **Fig 3**, we noticed a statistically significant 3-fold decrease in the circulatory cytotoxic CD8+ T cell abundance in atherosclerotic Pfn1^+/−(EC)^ vs Pfn1^+/+^ mice. A similar trend of 5.5-fold decrease was also observed in the aortic CD8+ T cell content in Pfn1^+/−(EC)^ atherosclerotic mice although the difference did not reach statistical significance owing to large animal-to-animal variability. Given that this trend was not observed in either splenic or bone marrow CD8+ T cell population, it is likely that CD8+ T cell mobilization from lymphoid organs rather than their development was negatively impacted in atherosclerotic Pfn1^+/−(EC)^ mice. It is important to note that non-atherosclerotic mice did not show any effect of partial loss of EC Pfn1 on either circulatory or aortic CD8+ T cell abundance (**Fig S2**). Interestingly, when analyzed for the relative content of B cells between the two groups of animals, both splenic and aortic B cell content in atherosclerotic Pfn1^+/−(EC)^ mice showed signs of elevation (by ∼3.5 fold) relative to Pfn1^+/+^ animals although the difference did not reach statistical significance because of large animal-to-animal variability (**Fig 3**). Somewhat consistent with these findings, non-atherosclerotic Pfn1^+/−(EC)^ mice also exhibited a sign of elevated circulating B cells relative to Pfn1^+/+^ mice with the p-value close to being significant (p=0.07); however, a reverse trend was found when splenic B cell content was compared between the two groups of animals in non-atherosclerosis setting (**Fig S2**). We recently showed that complete loss of vascular EC Pfn1 induces a selective bias of myeloid-derived cells (monocytes/macrophages) toward a pro-inflammatory phenotype *in vivo* (17)]. Consistent with these findings, anti-inflammatory Ly6C^lo^ monocytes were present at a lower level in atherosclerotic bone marrow and non-atherosclerotic spleen isolates of Pfn1^+/−(EC)^ relative to Pfn1^+/+^ mice; however, these changes were not preserved in the aortic immune cell compositional analyses (**Fig 2**). In summary, these immunological profiling studies demonstrate that the most noteworthy changes in the immune cell composition of the aorta (the main pathologic tissue of interest) elicited by partial loss of EC Pfn1 in atherosclerosis setting are trends of substantially diminished and elevated CD8+ T cells and B cells, respectively. Interestingly, these immunological trends are very different from those exhibited by mice completely deficient in endothelial Pfn1(17). Therefore, immunological consequences of Pfn1 depletion in ECs may be complex and cannot be simply explained by the degree of loss of expression in a dose-dependent manner.

**Figure 3.**
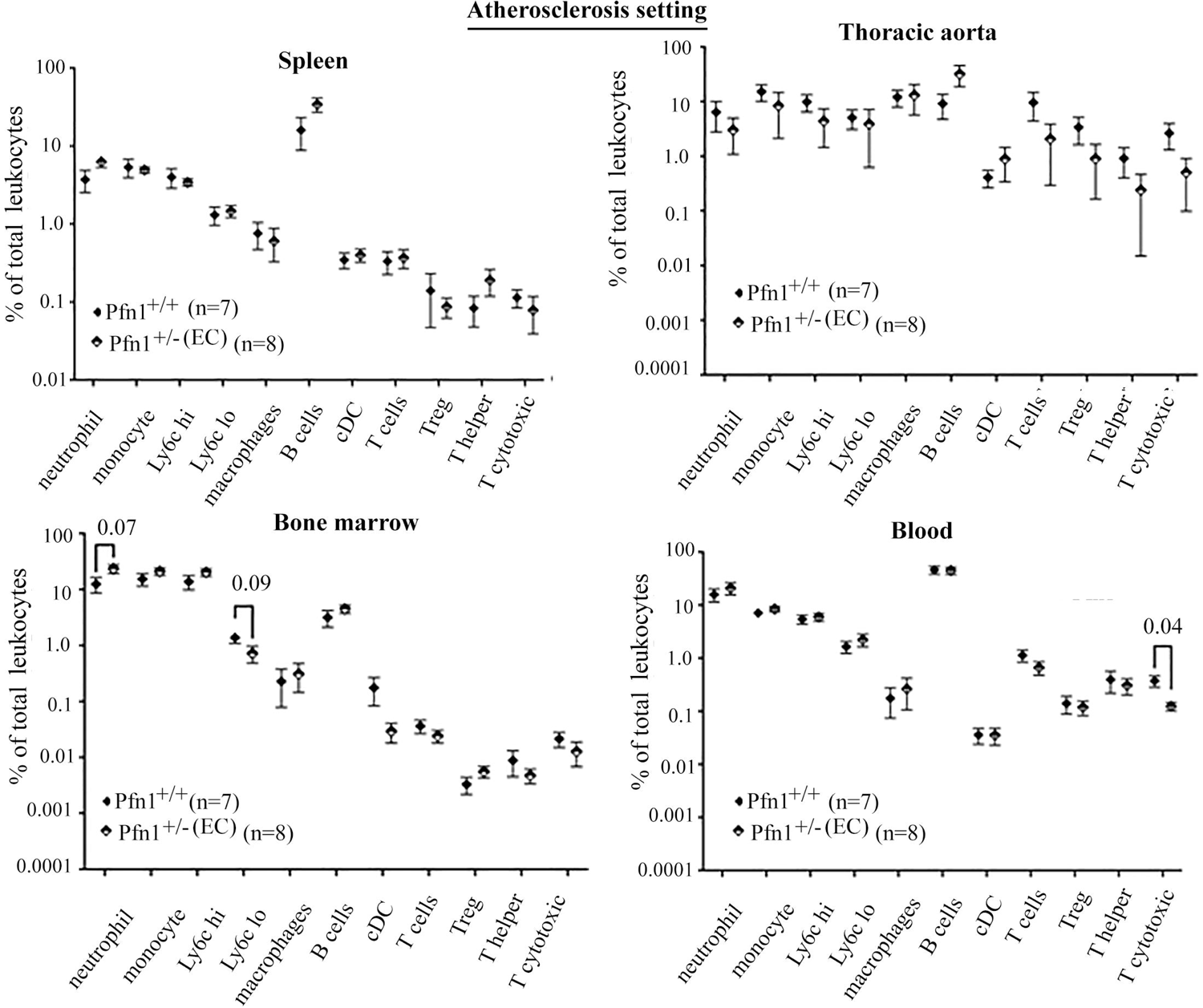
Immuneprofiling of athersclerotic Pfn1^+/+^ and Pfn1^+/−(EC)^ mice. Flow cytometry-based analyses of the relative abundance of various immune cell subtypes (as % of total leukocytes) in the circulation (whole blood) and various organs/tisssues (spleen, aorta and bone marrow) of atherosclerotic Pfn1^+/+^ and Pfn1^+/−(EC)^ (‘n’ indicates the number of animals in each group pooled from mutiple experiments; ‘p’ values when significant or close to being significant are indicated).

To gain further mechanistic insights into how partial loss of EC Pfn1 may alter circulating/aortic composition of select immune cell sub-types in atherosclerosis setting, we performed a broad assessment of circulating cytokine/chemokines in the two groups of animals utilizing a Luminex assay probing for a panel of 32 immunomodulatory factors. As per these analyses, IP-10/CXCL10 and IL7 are the only two immunomodulatory factors that were differentially abundant within statistical significance with the average circulating CXCL10 and IL7 levels being ∼40% and ∼70% lower, respectively, in Pfn1^+/−(EC)^ relative to Pfn1^+/+^ atherosclerotic animals (**Figs 4A-B**). Similar Luminex analyses performed with serum of non-atherosclerotic animals did not reveal any significant difference in circulating level of either CXCL10 or IL7 between the two groups, suggesting that the effect of partial loss of EC Pfn1 on these cytokines is specific to atherosclerosis setting (**Figs 4A-B**). CXCL10 is associated with pro-inflammatory immune cell activation, specifically playing a prominent role T cell recruitment in chronic inflammation (31,32). Genetic ablation of CXCL10 leads to reduced CD8+ T cell accumulation and enhanced Treg activity resulting in a prominent two-fold reduction in atherosclerotic lesion formation in ApoE^−/−^ mice(31). Reduced circulating CXCL10 level in Pfn1^+/−(EC)^ relative to Pfn1^+/+^ animals specifically in atherosclerosis setting may partly explain diminished circulating and aortic CD8+ T cell abundance as a consequence of partial loss of EC Pfn1. Interestingly, these CXCL10-associated data diverges from our recently published findings of prominent CXCL10 elevation in response to complete loss of EC Pfn1 *in vitro* and *in vivo* (17). IL7, a cytokine that triggers Th1 and Th17 response, has pro-atherogenic function (33,34). Microtubules play an important role in facilitating cytokine transport and release (35). Although in addition to regulating actin dynamics, Pfn1 promotes microtubule growth (36), we only saw a marginal 10% reduction in microtubule content in Pfn1-haploinsufficient EC relative to wild-type cells. Furthermore, given that out of 32 immunomodulatory factors that we probed, we only saw changes in the circulating levels of CXCL10 and IL7, we speculate that Pfn1-dependent changes in the circulating levels of CXCL10 and IL7 are likely due to alteration in specific signaling pathways rather than caused by subtle changes in the microtubule dynamics, if any. In hypertension mouse model, Pfn1 overexpression has been shown to result in elevated p38-MAPK activation (37). Given prior evidence for p38-MAPK activation linked to induction of CXCL10 expression (38), Pfn1-dependent changes in p38-MAPK activation could be a potential contributing factor in dampening of at least CXCL10 in Pfn1^+/−(EC)^ mouse model in our study. Since p38-MAPK activation is elevated in inflammatory conditions including atherosclerosis (39), it might also explain why we detected suppression of CXCL10 induced by partial depletion of Pfn1 in atherosclerotic mice only.

**Figure 4.**
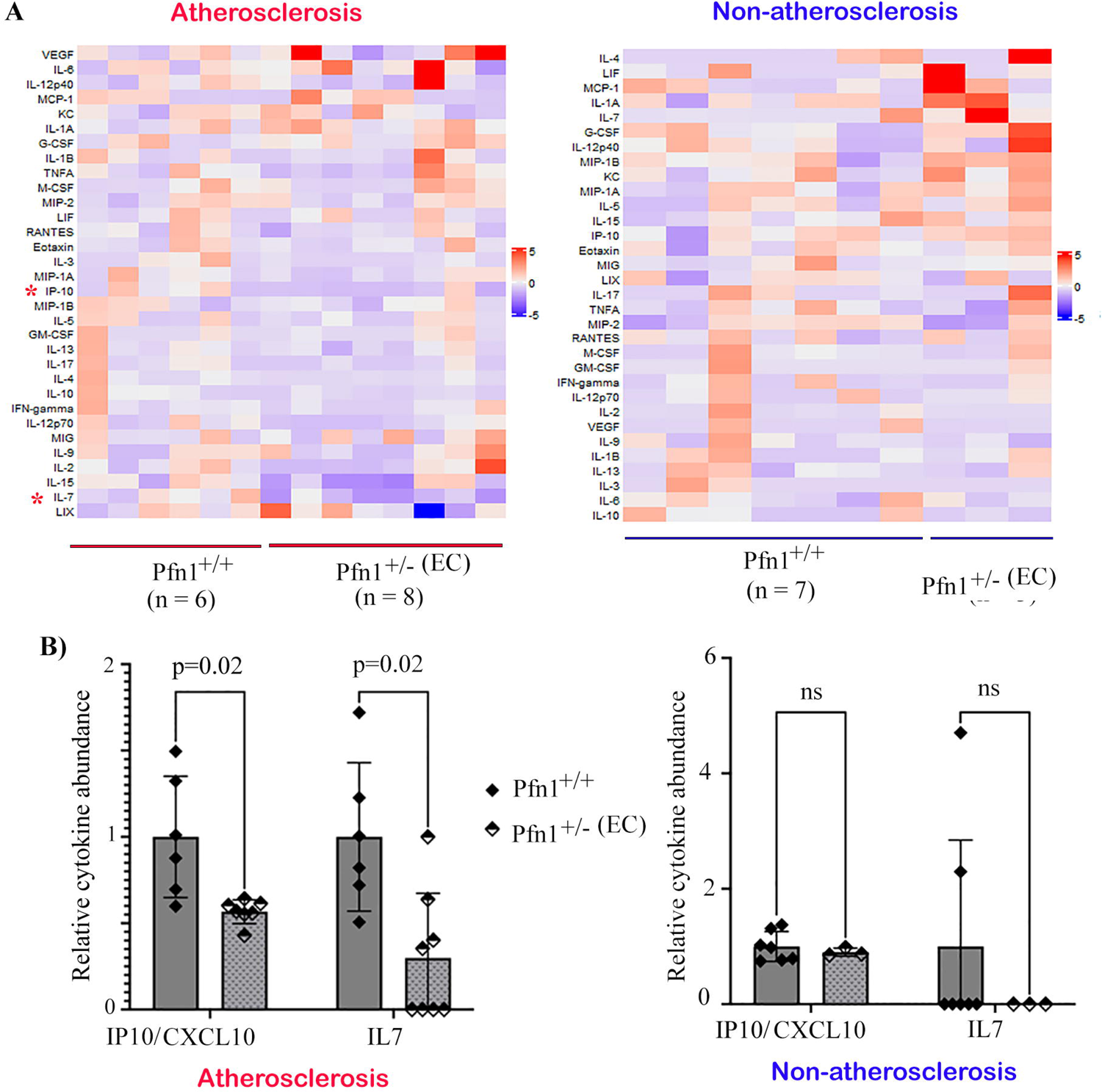
Luminex based profiling of circulating cytokines and chemokines of atherosclerotic vs non-atherosclerotic Pfn1^+/+^ and Pfn1^+/−(EC)^ mice. **A-B)** Heat-plots showing relative abundance of indicated cytokines and chemokines in the serum of atherosclerotic (*panel A*) and non-atherosclerotic (*panel B*) mice of indicated genotypes (data for each analyte normalized to the corresponding mean value calculated of Pfn1^+/+^ mice; immunomodulatory factor(s) that are differentially abundant within statistical significance are indicated by asterisk. **B)** Relative serum abundance of CXCL10 and IL7 between the two groups of animals in atherosclerosis vs non-atherosclerosis settings (‘n’ indicates the number of animals in each group pooled from mutiple experiments; ns – not significant).

In conclusion, this study for the first time investigates the impact of cell-type-specific partial loss of Pfn1 expression on atherosclerosis development. Although changes in EC play a key role in the initiation of atherosclerosis, our studies show that partial loss of Pfn1 expression selectively in ECs is not sufficient to confer atheroprotection *in vivo*. However, we believe that a substantial >3-fold reduction in the cytotoxic T cell abundance in the circulation as well as the aorta (a finding consistent with robust dampening of select pro-atherogenic cytokines namely, CXCL10 and IL7) in atherosclerotic Pfn1^+/−(EC)^ mice is physiologically meaningful. Given preclinical and human correlation studies supporting a pro-atherosclerotic role of CD8+ T cells (25–27), partial depletion of endothelial Pfn1 could still have beneficial effects, in particular, against initiation of the disease. Endothelial Pfn1is also critical for neovascularization(18). Although not studied here, given neovascularization in the atherosclerotic plaques promotes plaque growth and instability (40), partial depletion of EC Pfn1 may also have additional beneficial effects in the advanced stage of the disease through attenuating angiogenesis. Nonetheless, our results suggest that previously reported atheroprotective phenotype of global Pfn1^+/−^ mice (16) is attributed to combined effects of Pfn1 depletion in additional cell types that are relevant for atherosclerosis progression. In this context, it is particularly important to note that in atherosclerotic plaques, Pfn1 abundance is elevated in both intracellular and extracellular spaces(14). Although Pfn1 is best known for its intracellular functions, Pfn1 is also externalized by various cells (41) and cell culture experiments show that perturbing intracellular Pfn1 content in EC commensurately alters the externalized Pfn1 content(42). Interestingly, soluble Pfn1 is capable of stimulating vascular smooth muscle cell (SMC) migration and proliferation (important features of atherosclerosis) (14).Therefore, it is possible that cell-secreted Pfn1 can have additional pro-atherogenic effect through an extracellular action on vascular SMC. In that case, one would expect more substantial reduction in the extracellular Pfn1 content in the vascular intima when Pfn1 is globally reduced in all cell types vs in selected cell type. This could partly explain the lack of atheroprotective phenotype when Pfn1 is solely depleted in ECs. Additionally, immune infiltration of the vascular space and plaque formation/growth can be also affected when Pfn1 is additionally depleted in either CD8 T cells or those of myeloid origins (macrophages, monocytes), features that are not replicated in our *in vivo* model. Therefore, it will be important to investigate in the future whether attenuated Pfn1 expression in vascular SMC and/or sub-types of immune cells offer additional benefits that could potentially explain atheroprotection conferred by global Pfn1 haplo-insufficiency.

## ACKNOWLEDGMENTS

The authors wish to acknowledge the contribution of Pooja Chawla in the Roy lab for her technical assistance. AAG was supported by a NHLBI grant F31HL160118-01, ARCS funding and a Cardiovascular Bioengineering T32 training grant from the NHLBI (5T32 HL 76124-14). DG was supported by NCI K99-CA267180 grant, a National Cancer Center fellowship, and an Imaging Sciences in Translational Cardiovascular Research T32 training grant from NHLBI (T32-HL 129964). Research in the Roy lab has been supported by grants from NCI (R01CA248873, R01CA271095), Department of Defense (W81XWH-19-1-0768), and NEI (R21EY032632).

