## Supplementary data for "Haplo-insufficiency of Profilin1 in vascular endothelial cells is beneficial but not sufficient to confer protection against experimentally induced atherosclerosis"

**Allen-Gondringer et al. Fig S1**

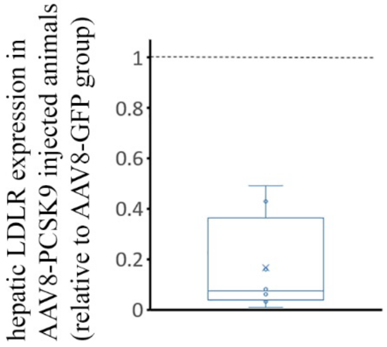


**Figure S1**: **Confirmation of PCSK9 AAV-driven suppression of LDLR expression in mouse liver**. Box-and-whisker plot displaying hepatic LDLR expression in AAV8-PCSK9 injected mice (n=8; pooled from $Pfn1^{+/+}$ and $Pfn1^{+/-(EC)}$ mice) relative to that in control AAV8-GFP injected $Pfn1^{+/+}$ mice (n=3).

**Allen-Gondringer et al. Fig S2**

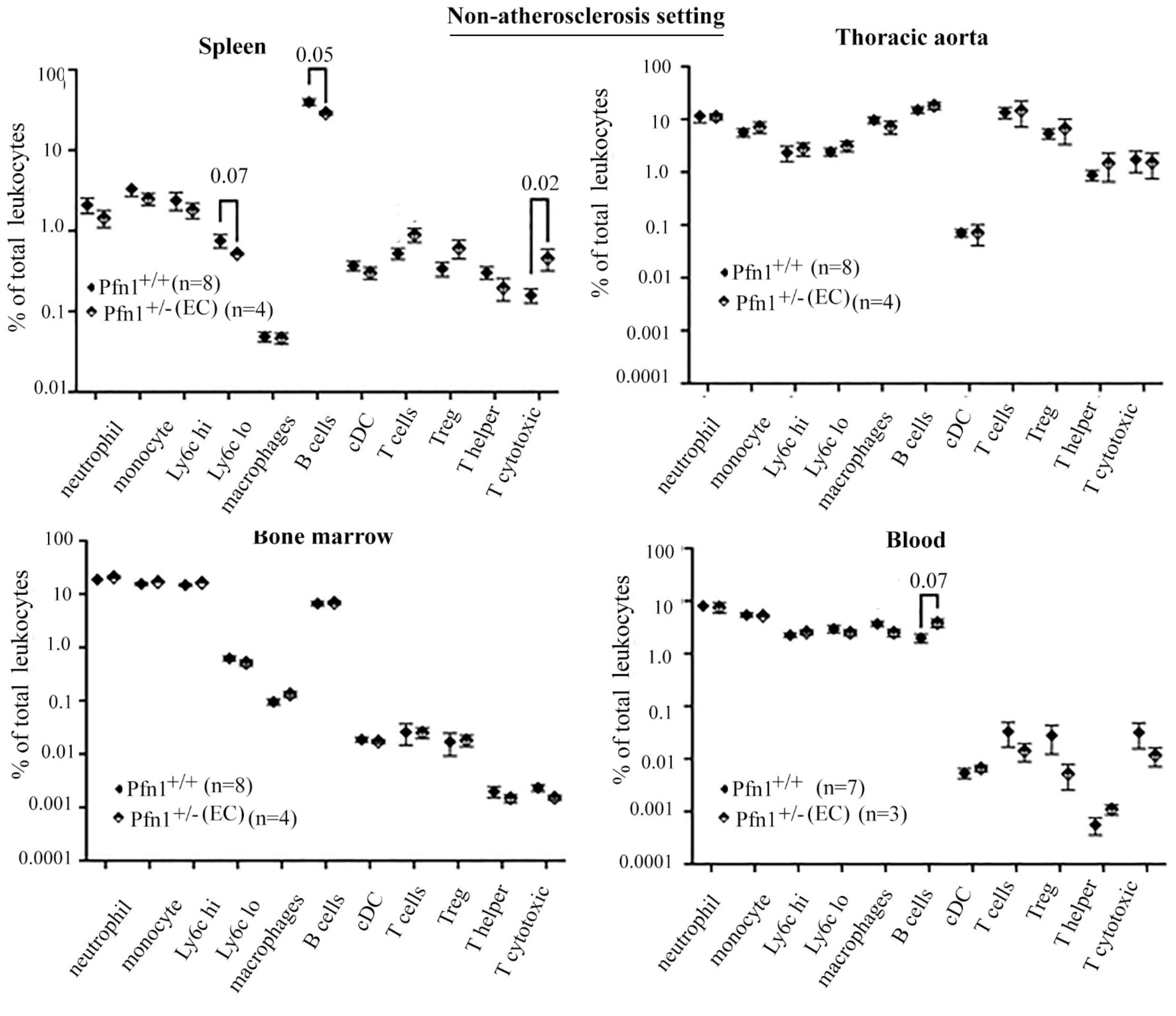


**Figure S2**. **Immuneprofiling of non-athersclerotic Pfn1$^{+/+}$ and Pfn1$^{+/-(EC)}$ mice**. Flow cytometry-based analyses of the relative abundance of various immune cell subtypes (as % of total leukocytes) in the circulation (whole blood) and various organs/tisssues (spleen, aorta and bone marrow) of non-atherosclerotic Pfn1$^{+/+}$ and Pfn1$^{+/-(EC)}$ ('n' indicates the number of animals in each group pooled from mutiple experiments; 'p' values when significant or close to being significant are indicated).
